# Rapid, Non-Destructive Visualization of α-Zein Expression and Grain Protein Concentration in Maize Using the *Floury2*-RFP Reporter Transgene

**DOI:** 10.64898/2026.05.05.723001

**Authors:** Catherine Li, Nicholas J. Heller, Christine Tiskevich, Stephen P. Moose

**Author notes:** **Correspondence** Stephen Moose.

## Abstract

Kernel composition traits in maize, including protein accumulation, are of broad interest. The amount of the most abundant proteins in maize endosperm, the α-zeins, can vary dramatically among genotypes and in response to soil nitrogen supply. Targeted reductions in α-zein accumulation can improve nitrogen utilization and the nutritional quality of maize grain but have traditionally required expensive and destructive phenotyping methods. The *Floury2*-RFP (*Fl2*-RFP) reporter gene enables rapid, non-destructive visualization of α-zein accumulation in individual maize kernels under white light. This feature is due to the high expression level programmed by the *Fl2* promoter, the stability of zein proteins, and the use of monomeric RFP, which emits fluorescence without the need for multimerization. This study aimed to develop a method to quickly document and quantify *Fl2*-RFP accumulation using camera or smartphone images of either ears or shelled kernels. Results show images of shelled kernels processed with FIJI software capture the *Fl2*-RFP reporter phenotype better than images of ears. *Fl2*-RFP confirms the strong maternal control of α-zein accumulation and, like grain protein concentration, responds to soil nitrogen supply. The *Fl2*-RFP phenotyping pipeline effectively quantified *Fl2*-RFP accumulation by color features from both camera and smartphone images. Smartphone imaging of *Fl2*-RFP in a diverse population of inbreds followed by elastic net regression of extracted image features predicted kernel protein concentration, as measured by near-infrared spectroscopy, with moderate accuracy (*R*^2^ = 0.68, MAE = 0.76, RMSE = 0.93). The spectral features that were most predictive of kernel protein concentration varied depending on whether the background endosperm color was white or yellow. The integrated analysis of *Fl2*-RFP intensity and grain protein concentration indicates genetic variation for kernel protein accumulation and N-responsiveness that is distinct from the well-studied α-zeins. Our findings highlight the *Fl2*-RFP reporter gene as a valuable tool for investigating the genetic complexity of grain protein concentration and associated traits in maize.

## 1 Introduction

Maize (*Zea mays* L.) is a globally important cereal crop, due to its high yield and diversity of uses in food, feed, fuel, and industrial products. For each use case, a different seed composition is desirable. The nutritional and industrial value of maize is fundamentally determined by the composition of its seeds, which are structurally divided into the pericarp, endosperm, and germ (embryo). Whereas the germ comprises about 10% of the dry weight in a maize kernel, and the pericarp 5%, the endosperm constitutes much of the remaining 85% and contains 80% of the seed protein (Tollenaar and Dwyer, 1999).

The Illinois Long Term Selection Experiment (ILTSE) was initiated in 1896 at the University of Illinois by Cyril G. Hopkins to alter grain composition traits through selective breeding (Hopkins, 1899). Commercially relevant modern maize lines typically have a grain protein concentration of 8– 15% (Manthey, 2016), whereas the ILTSE has since produced populations with the known phenotypic extremes for grain protein concentration: the Illinois High Protein (IHP) with 30% protein and Illinois Low Protein (ILP) at 4%. The Illinois Reverse High Protein (IRHP) and Illinois Reverse Low Protein (IRLP) populations fall intermediate between the two. Collectively, these populations are known as the Illinois Protein Strains (IPS; Moose et al., 2004).

Zeins, also known broadly as prolamins across cereal species, are the most abundant seed storage protein in maize. Zeins constitute roughly half of the endosperm protein, and thus nearly half of the total seed protein. The dominance of zeins in the endosperm is the primary reason for the poor protein quality of maize, as zeins are deficient in the essential amino acids lysine, methionine, and tryptophan (Paulis and Wall, 1977). Accordingly, most attempts to improve the nutritional quality of maize protein involve altering zein content (Flint-Garcia et al., 2009). Of the zein subfamilies, the α-zeins are the largest (Coleman and Larkins, 1999). Biochemical analysis of the IPS throughout the experiment revealed that changes in grain protein concentration resulted primarily from altered accumulation of the 19- and 22-kD α-zeins, with greater amount of zeins in IHP endosperm compared to ILP (Bhattramakki et al., 1996).

The *Floury2*-RFP (*Fl2*-RFP) reporter line was created by the collaborative efforts of the Maize Cell Genomics project as one of many fluorescent protein reporter lines (Mohanty et al., 2009; Wu et al., 2013; Krishnakumar et al., 2015). The reporter line consists of a monomeric Red Fluorescent Protein (RFP) derived from the coral *Discosoma* genus (Campbell et al., 2002) fused to the *Floury2* (*Fl2*) gene, which encodes a 22-kD α-zein. *Floury2* is the second highest expressed 22-kDa-zein in the inbred variety B73, accounting for ∼20% of the expression (Song and Messing, 2003; Feng et al., 2009). Due to the high expression of the relatively stable *Floury2* gene, and the fact that monomeric RFP does not require multimerization to emit fluorescence, the *Fl2*-RFP phenotype can be visualized in white light without specialized equipment (Lucas et al., 2013). When introgressed into inbred lines derived from the Illinois Protein Strains and B73, the mRNA expression and FL2-RFP accumulation in these transgenic lines appear to follow the spatial and temporal patterns of the native *Floury2* gene (Woo et al., 2001), demonstrating the *Fl2*-RFP transgene is a valuable tool for tracking variation in seed protein concentration (Lucas et al., 2013).

Analytical methods for protein quantification in maize kernels include traditional wet chemistry (Kjeldahl, Dumas), and advanced spectroscopic techniques such as Nuclear Magnetic Resonance (NMR) and Near Infrared Reflectance (NIR) (Dudley and Lambert, 2004; Moose et al., 2004). NIR has become the method of choice for rapid, high-throughput screening in breeding programs, offering accuracy and reproducibility when combined with appropriate calibration models. However, these methods require specialized equipment and measure all amide bonds and proteins in the sample. The possibility to specifically track inheritance of the α-zeins would allow for a more targeted approach to dissecting the genetic control of seed protein concentration and composition.

Many traditional phenotyping approaches are reliant on manual measurements and visual scoring, which in turn are labor-intensive, subjective, and inherently limited in scale and resolution. In the past decade, the use of image-based phenotyping techniques has expanded rapidly, enabling high-throughput and reproducible extraction of maize kernel and ear traits with digital cameras and open-source software (Miller et al., 2017, Makanza et al., 2018, Warman et al., 2021). These phenotyping platforms can quantify biologically relevant features from both intact ears and individual kernels, though most existing pipelines focus on morphological traits, such as kernel length, width, and shape, as they can be used to estimate yield components (Miller et al., 2017, Makanza et al., 2018). Warman et al. (2021) further demonstrated the utility of combining fluorescence-based imaging with these methods by using Green Fluorescent Protein (GFP) to identify and count kernels, highlighting the potential of reporter systems for trait-specific phenotyping. As the intensity of the *Fl2*-RFP phenotype varies with different genetic backgrounds and environmental conditions, quantifying its expression allows us to more easily characterize the regulation of an individual α-zein gene. The methods presented here enable accurate, repeatable, and cost-effective image-based quantification of the *Fl2*-RFP reporter.

## 2 Materials and methods

### 2.1 Plant Materials

The IHP1, ILP1 IRHP1 and IRLP1 inbred lines derived from the Illinois Protein Strains have been described previously (Uribelarrea et al., 2004). The *Fl2*-RFP transgene was introgressed into each of these inbred lines and the inbred line B73 by at least six generations of backcrossing followed by selfing (Lucas et al., 2013). Reciprocal crosses were created by crossing one parent heterozygous for the *Fl2*-RFP transgene with the other parent lacking the transgene, which produced grain with a broad range of protein concentrations and FL2-RFP intensities. Hybrids were produced by crossing the B73 inbred carrying the *Fl2-RFP* transgene as female with either ILP1, IHP1 or IRHP1. The population of diverse inbreds expressing FL2-RFP was created by crossing either IHP1 or ILP1 carrying the *Fl2*-RFP transgene with an elite hybrid (PHG39 x PHJ31) and self-pollinating FL2-RFP-expressing kernels for six generations while selecting for a range of visual FL2-RFP intensity.

The *Fl2*-RFP introgressions into the IPS inbreds and B73, as well as their derived hybrids, were grown at the Crop Sciences Research and Education Center in Urbana, IL in 2012, 2013, and 2024. The population of diverse FL2-RFP inbreds was grown at the same location in 2019. Each genotype was grown in replicated single row plots 5.34m in length with 76.2-cm row spacing. Hybrids were grown in a split-plot design, where the same hybrid was grown in paired plots receiving either no fertilizer or nitrogen fertilizer in the form of granular ammonium sulfate applied at a rate of 200 kg N ha^−1^, when the plants reached the V3 growth stage.

### 2.2 Kernel Microscopy

Three-dimensional images were collected on a DeltaVision deconvolution microscope using a 60X lens and 0.2 μm Z-step optical sections, as described in Bass et al. (2014). Nuclei were isolated from endosperm cells at 14 days after pollination (DAP) and stained with DAPI. Datasets of DAPI and Red Fluorescence were captured for entire nuclei and deblurred by deconvolution. Images were adjusted by linear scaling of intensity shown as single wavelength or pseudocolored image overlay of both channels (DAPI and CY3).

Images of 24 DAP IHP1 and ILP1 kernels were taken using the Zeiss confocal laser scanning microscope (LSM 710) with a 40x objective. A feature of this microscope is the ability to combine aspects of light and fluorescence microscopy, which allows for the visualization of the different cell layers, including the pericarp, the aleurone and the subaleurone, along with the visualization of *Fl2*-RFP expression within the different cell layers. From these images and others like them, it is possible to quantify the number of cellular layers expressing the *Fl2*-RFP transgene.

### 2.3 Image Acquisition & RFP Phenotyping

At maturity, the ear is harvested and immediately wrapped in a paper bag to avoid photo-bleaching. The Fl2-RFP phenotype is visualized with white light; thus any number of image-capturing devices can be used to directly record and digitally preserve the phenotype, and the photographs can be analyzed with computer software. If desired, a photograph of the intact ear is captured before shelling. In lines that are still segregating for RFP, kernels containing the *Fl2*-RFP reporter are subsetted for image analysis. The shelled RFP kernels from those ears were laid out embryo-side down and non-overlapping on a solid, contrasting colored background. Photographs were taken in a room with controlled lighting and no windows.

In this study, a Canon EOS 5D was used to capture images of the intact hybrid ears. For all pictures of shelled kernels, a Nikon D60 or iPhone 13 mini was used. To prepare the images for analysis, a custom Adobe Lightroom preset developed with the SpyderCheckr® 24 color card (datacolor, Lawrenceville, New Jersey, United States) is applied to color correct the photo. If needed, Adobe Lightroom AI masking is used to blackout the background. Then, an adapted macro from Strock (2021) is deployed in FIJI (Schindelin et al., 2012) to automate the remainder of the process. This macro uses thresholding to separate seeds from background, clear the background so that only the seeds remain, duplicates and separates the image into RGB, LAB, and HSB color spaces, and records the mean, max, min, mode, and standard deviation for each channel. Shape descriptors including area, aspect ratio, circularity, roundness, and solidity of each kernel are also recorded. The thresholded and masked images are saved for QC. This macro automates the process and takes ∼5 seconds per image.

Grain protein of individual ears was measured with a Near Infrared Analyzer (PerkinElmer Perten DA 7200) equipped with a custom calibration to handle the extremes in maize grain composition from the ILTSE and variation in endosperm color.

### 2.4 Analysis

For a preliminary assessment of the data, Pearson correlation analysis was used to evaluate how correlated the extracted features were between the imaging methods, and to total grain protein concentration. Due to the high dimensionality, multicollinearity, and low sample number in this dataset, elastic net regression (Zou and Hastie, 2005) implemented in R (v4.5.3; R Core Team 2026) using glmnet (v4.1.10; Tay et al. 2023) was used for feature selection and prediction with alpha = 0.15, leaning closer to ridge regression, and leave-one-out cross-validation. Kernel endosperm color was imputed as a dummy variable, with 0 = white and 1 = yellow. Model performance was evaluated using the coefficient of determination (R^2^), mean absolute error (MAE), and root mean square error (RMSE).

## 3 Results

### 3.1 Visualization of the *Fl2*-RFP reporter phenotype at different scales

The *Fl2*-RFP reporter transgene (Figure 1A) consists of RFP inserted in-frame to the native genomic sequence of *Floury2* and expressed as a fusion to the translated protein (Wu et al., 2013; Krishnakumar et al., 2015). This includes 3042-bp of sequence upstream of the translational start site and 1082-bp downstream of the α-zein stop codon, where mRFP1 was fused in-frame near the C-terminus, but keeping the last 16 amino acids of the zein coding sequence to preserve functions important for interactions with other zeins and aggregation into protein bodies (Kim et al., 2002). Inclusion of the native regulatory sequences facilitates the preservation of proper tissue-specific and temporal expression, providing insight into regulatory elements that act at the level of transcription as well as those involved in the processing, localization, and turnover of α-zein mRNA and protein.

**Figure 1.**
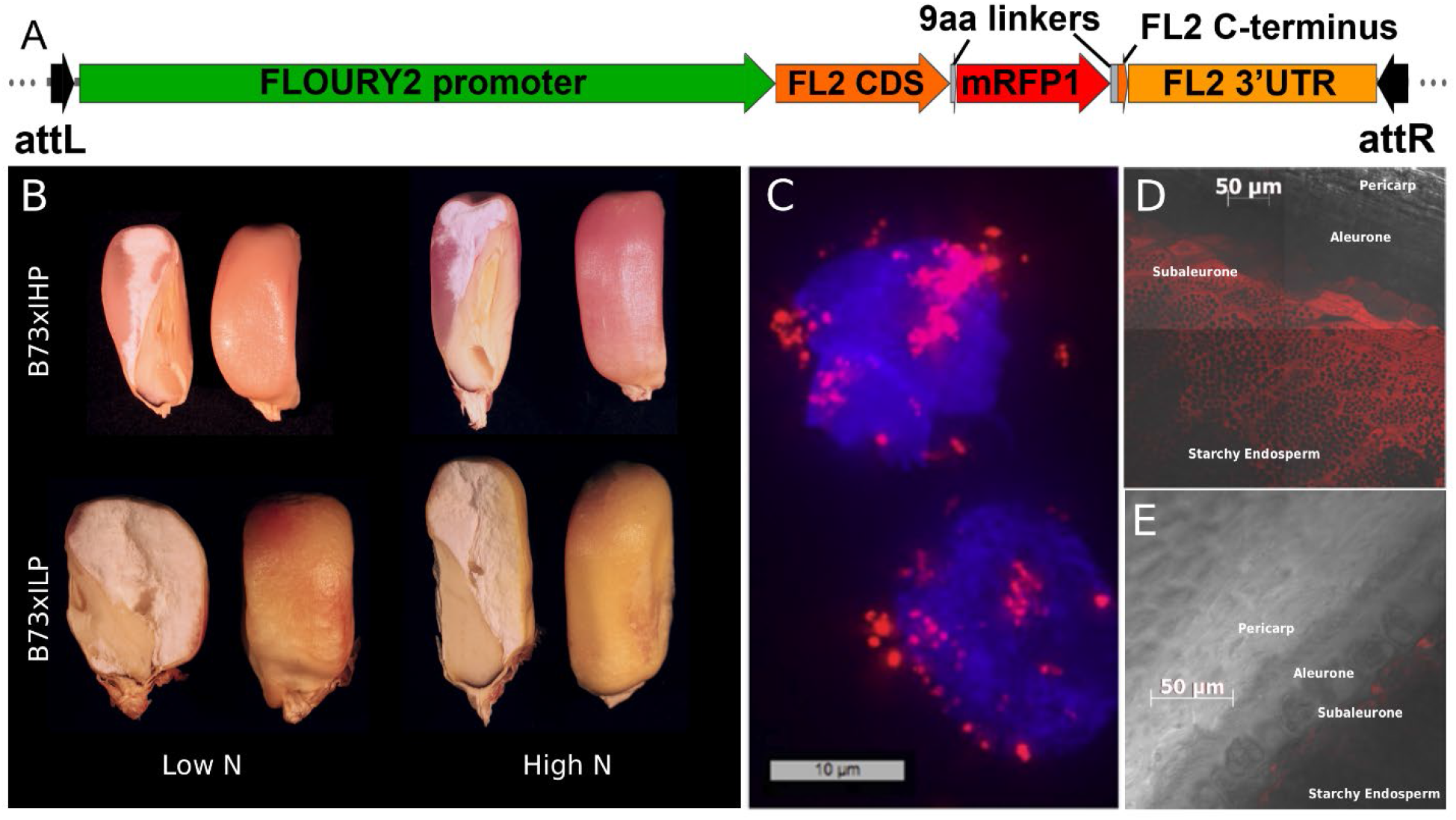
Spatial accumulation of *Fl2*-RFP within maize kernels. **(A)** Cartoon of FLOURY2 (*Fl2)*-RFP transgene structure. **(B)** Cross-sections of representative kernels from IPS hybrid ears containing the FL2-RFP transgene **(C)** 3D deconvolution microscopy of two nuclei isolated from 14 DAP kernels containing FL2-RFP. The blue DAPI signal stains nuclear chromatin, with the FL2-RFP fusion protein accumulating as red signals. **(D)** Detection of FL2-RFP fluorescence using light and fluorescence confocal microscopy of endosperm cells imaged at 24 DAP from IHP1 kernels and **(E)** from ILP1 kernels.

We previously demonstrated that the expression profile of the *Fl2*-RFP reporter closely mirrors that of the entire 22-kDa α-zein subfamily, and even correlates with total grain protein concentration in the IPS, where variation in α-zein content is under strong genetic control (Lucas et al., 2013). Representative images of ears from the *Fl2-RFP* transgene introgressed into the IPS or B73 inbreds, or crosses between these inbreds, are shown in Figure S1. FL2-RFP intensity is weakest in ILP1, progressively increasing in IRHP1, IRLP1 and IHP1. FL2-RFP intensity in B73 appears visually similar to IRHP1. Ears from reciprocal crosses show a strong maternal effect on FL2-RFP expression. After planting seeds from the B73: RFP x ILP1 and B73: RFP x IHP1 hybrids (Fig. S1D) in field trials and self-pollination of the ears, representative images of mature kernels illustrate that both genetic background and environmental nitrogen availability influence the intensity and spatial distribution of the reporter phenotype (Figure 1B). FL2-RFP accumulation is readily visible in white light and reflects the pattern expected for the native FLOURY2 protein: high in the subaleurone and vitreous endosperm, and low in the starchy regions (Woo et al., 2001). Also evident is the long-known relationship between reduced zeins and a floury/opaque kernel phenotype for the ILP hybrid kernels (Hopkins et al., 1903) compared to the vitreous appearance of IHP hybrid kernels.

Further confirmation that the accumulation of *Fl2*-RFP mimics that of endogenous α-zeins is obtained from its subcellular localization (Figure 1C). The bright foci of RFP appear tethered to the isolated nuclei, which are easily visualized by the blue DAPI staining of chromatin. These observations are consistent with the known synthesis of α-zeins in protein bodies from the rough endoplasmic reticulum associated with the nuclear envelope (Lending and Larkins, 1989). In both IHP1 (Figure 1D) and ILP1 (Figure 1E), *Fl2*-RFP expression is the strongest in subaleurone cells, with a progressive reduction in intensity into the starchy endosperm. These images show that *Fl2*-RFP expression extends much further into the kernel in IHP1 compared to ILP1. Greater than twenty cell layers of the subaleurone and starchy endosperm accumulated FL2-RFP in IHP1, compared to only approximately five layers of subaleurone cells in ILP1. The mRNA expression of α-zeins begins at 10 DAP and continues throughout grain fill (Lending and Larkins, 1989; Woo et al., 2001). *Fl2*-RFP expression could be detected as early as 12 DAP in IHP1 (Lucas et al. 2013). However, as ILP kernels accumulate very little zein, the FL2-RFP signal could only be detected at 24 DAP when using imaging settings that avoided saturating the much stronger signal present in IHP kernels. No expression is evident in the pericarp, aleurone or embryo, consistent with the exclusive accumulation of zeins in endosperm tissue (Lopes and Larkins, 1993; Woo et al., 2001).

### 3.2 The impacts of variation in genotype and nitrogen supply on FL2-RFP accumulation

Grain protein concentration shows strong maternal inheritance in the IPS, as well as other maize germplasm (East and Jones, 1920; Tsai, 1990). Grain protein concentration in hybrids derived from the ILTSE are generally intermediate between the parental means, and increases with higher nitrogen availability (Uribelarrea et al., 2004), except for ILP hybrids which remain largely unresponsive. This aspect of α-zein gene regulation is also evident in self-pollinated ears and kernels from the IPS hybrids containing *Fl2*-RFP (Figure 2A). Each hybrid ear is derived from crossing a white-endosperm and a yellow-endosperm parent (B73), therefore multiple kernel color combinations are observed. The known dramatic genetic differences for grain protein concentration among these three hybrids can be readily visualized by FL2-RFP intensity of unshelled ears. However, the impact of variable environmental nitrogen supply on FL2-RFP intensity is more subtle. While on the ear, the IHP hybrid ears grown with either low or high N appear visually similar in color, but once they are shelled, the kernels from IHP are visibly darker red under high nitrogen conditions. This difference is likely due to the spatial patterning of α-zeins, reflecting increased α-zein accumulation in peripheral endosperm tissues relative to the starchy crown.

**Figure 2.**
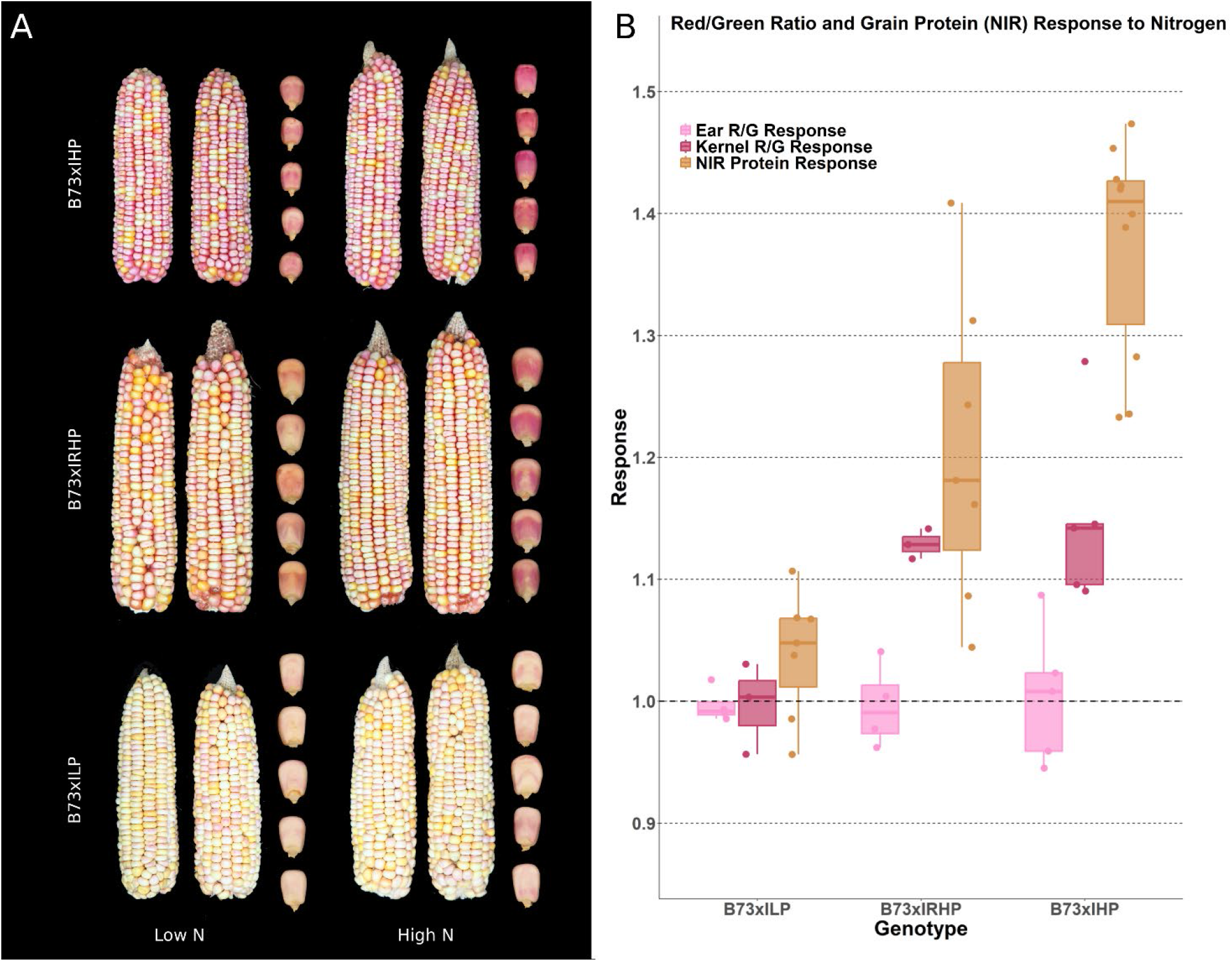
Genotype-by-nitrogen interaction for *Fl2*-RFP accumulation in maize hybrids. **(A)** Representative ears and kernels (enlarged to show detail) of IPS hybrids **(B)** Ears and kernels similar to those shown in Figure 1 were measured by NIR for grain protein concentration and for *Fl2-RFP* expression estimated by values of the ratio for RGB Red channel/RGB Green channel (R/G).

To better quantify the visually apparent genotype-by-nitrogen interaction for variation in Fl2-RFP and accumulation in maize kernels, we developed methods to measure Fl2-RFP intensity of shelled kernels from camera images. A visualization of this process is presented in Figure 3, which was then applied to kernels from ears similar to those shown in Figure 2A. Nitrogen responsiveness was defined as the ratio of the trait value when grown with high nitrogen to that under low nitrogen supply. Due to variability for either yellow or white endosperm, all subsequent analyses first separated kernels by endosperm color prior to imaging to reduce complexity. We did confirm that neither endosperm color nor *Fl2*-RFP presence/absence influenced grain protein concentration estimates using NIR. Among the many metrics extracted through the image phenotyping pipeline of the kernels, the ratio of mean RGB Red to Green (R/G) was the most correlated feature that also mirrored the pattern of grain protein concentration under different nitrogen conditions (*r* = 0.89 white kernels, *r* = 0.83 yellow kernels). This is due to R and G values both decreasing under high nitrogen conditions, which indicates that the kernel color is darker on those axes. As expected, the R/G ratio was highly correlated to CIELAB a* (*r* = 0.99); however, R/G better matches the nitrogen response pattern observed for grain protein concentration measured by NIR. Figure 2B shows that similar to direct visual assessment, the R/G metric did not change with N supply for any hybrid when measured on intact ears. The B73 x ILP hybrid showed limited nitrogen response grain protein concentration and no response for kernel R/G, whereas the B73 x IHP hybrid exhibited a strong positive response for both traits. The N response for grain protein concentration of the B73 x IRHP hybrid was intermediate between the ILP and IHP hybrids (Figure 2B), but the degree of N response for kernel R/G was similar to the IHP hybrid. Together, these results demonstrate that the *Fl2*-RFP reporter can provide a sensitive, image-based readout of both genetic and nitrogen-dependent variation in α-zein expression in maize kernels.

**Figure 3.**
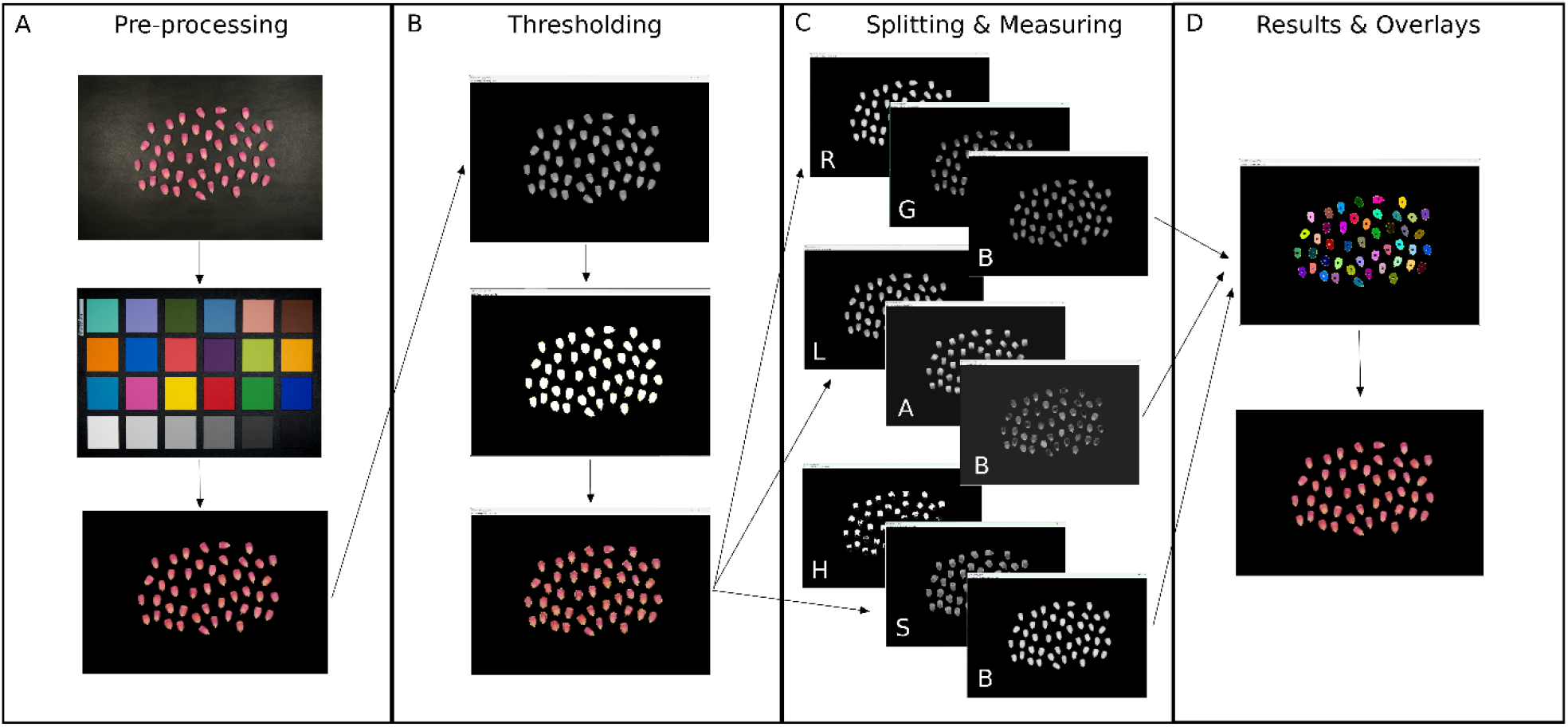
Visualization of image processing pipeline. **(A)** Original image, color balanced and blacked out background in adobe lightroom. **(B)** Once opened in FIJI, a mask is created using thresholding to separate the kernels from the background. **(C)** the original image is duplicated and split into different windows to measure each channel in RGB, LAB, and HSB colorspaces, along with shape descriptors. **(D)** the values for each kernel, along with an overlay mask image and thresholded image are saved for quality control.

### 3.3 Validation of the *Fl2*-RFP phenotyping pipeline in modern inbreds

To investigate the dynamics of the *Fl2*-RFP reporter in more modern germplasm, we developed a panel of inbred lines that vary for grain protein concentration and also express *Fl2-RFP*. Throughout the development of this panel, selection of FL2-RFP intensity was imposed to create a diversity of colors. The range of grain protein concentration in this population as measured by NIR was 8.64% to 15.42% (Figure 4A). Although the original hybrid was segregating for endosperm color due to its white- and yellow-endosperm parents, repeated self-pollination produced lines that are now uniform within each ear. The final panel consists of a total of 43 inbred lines, 16 with yellow endosperm, and 27 with white endosperm. Within this population, the visual relationship between FL2-RFP intensity and grain protein concentration is not always immediately apparent. Figures 4B and 4C illustrate two white-endosperm lines with similar protein, but distinct visual *Fl2*-RFP phenotypes. In contrast, Figures 4D and 4E show the more expected pattern in yellow-endosperm lines, where higher protein concentration corresponds to darker *Fl2*-RFP signal. These cases are just a few examples, as both endosperm color types produced ears with either strong or weak concordance between FL2-RFP intensity and grain protein concentration.

**Figure 4.**
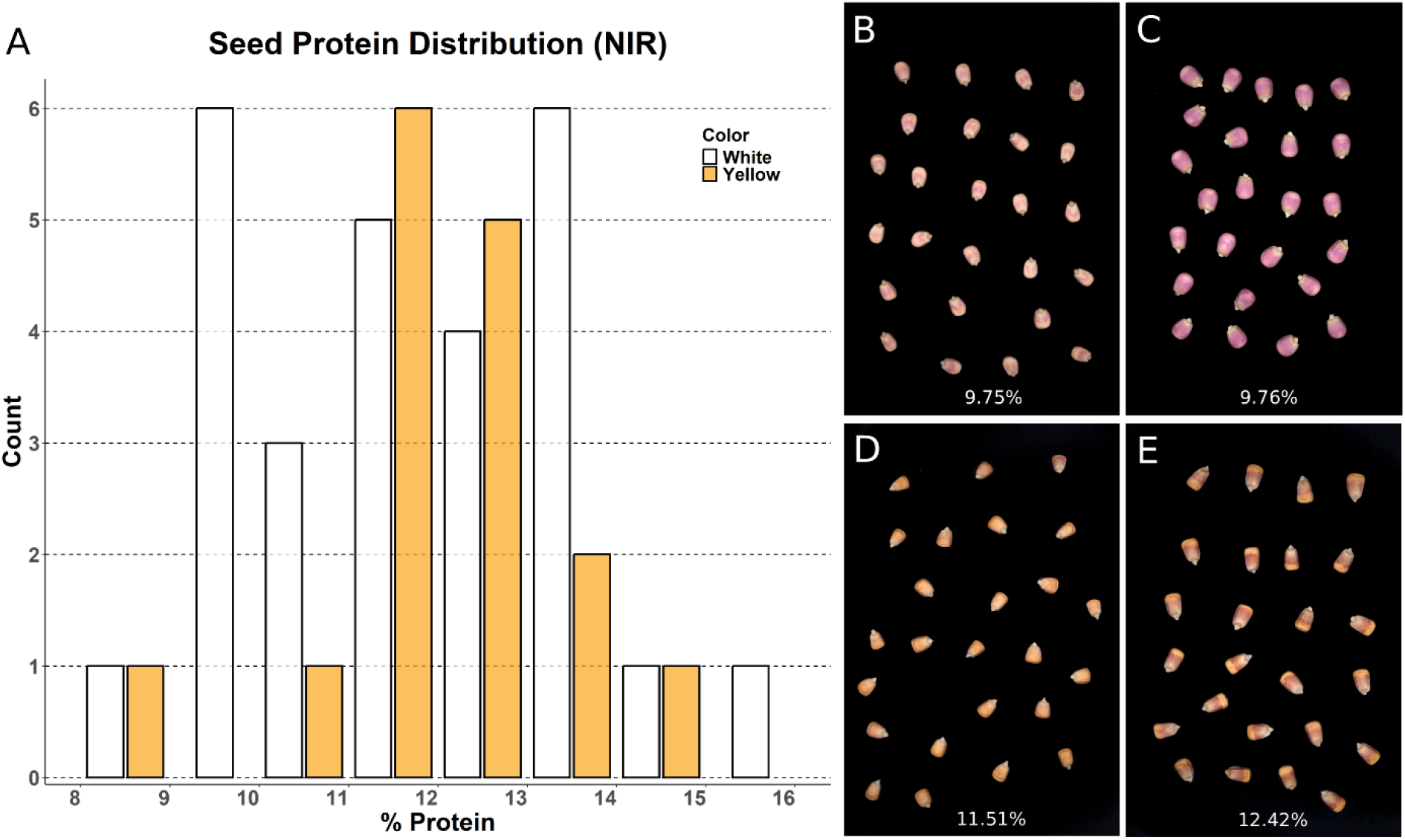
**(A)** Grain protein concentration distribution in a diverse inbred population carrying *Fl2*-RFP. **(B)** and **(C)** Example pictures of white kernels from this population with visual differences in *Fl2-RFP* reporter intensity but the same grain protein concentration. **(D)** and **(E)** yellow kernels where *Fl2*-RFP intensity is darker in kernels with higher grain protein concentration.

Furthermore, to test the effectiveness of different cameras in the phenotyping pipeline, pictures of the kernels were taken with both a digital camera and smartphone camera in succession. From these images, a total of 51 features were extracted, consisting of 5 shape descriptors and 46 color descriptors. In addition, endosperm color was manually annotated. We then evaluated cross-camera agreement and biological relevance by correlating the values of each predictor between devices and with grain protein concentration measured by NIR. The top 16 color traits that showed high concordance between cameras (*r* > 0.85) and their correlations to protein measured by NIR are shown in Figure 5. Not all color descriptors exhibited high camera concordance, indicating that the type of camera affects how some color features are captured, while others remain robust. By contrast, all shape descriptors were highly concordant across devices, reflecting their relative insensitivity to lighting and device differences.

**Figure 5.**
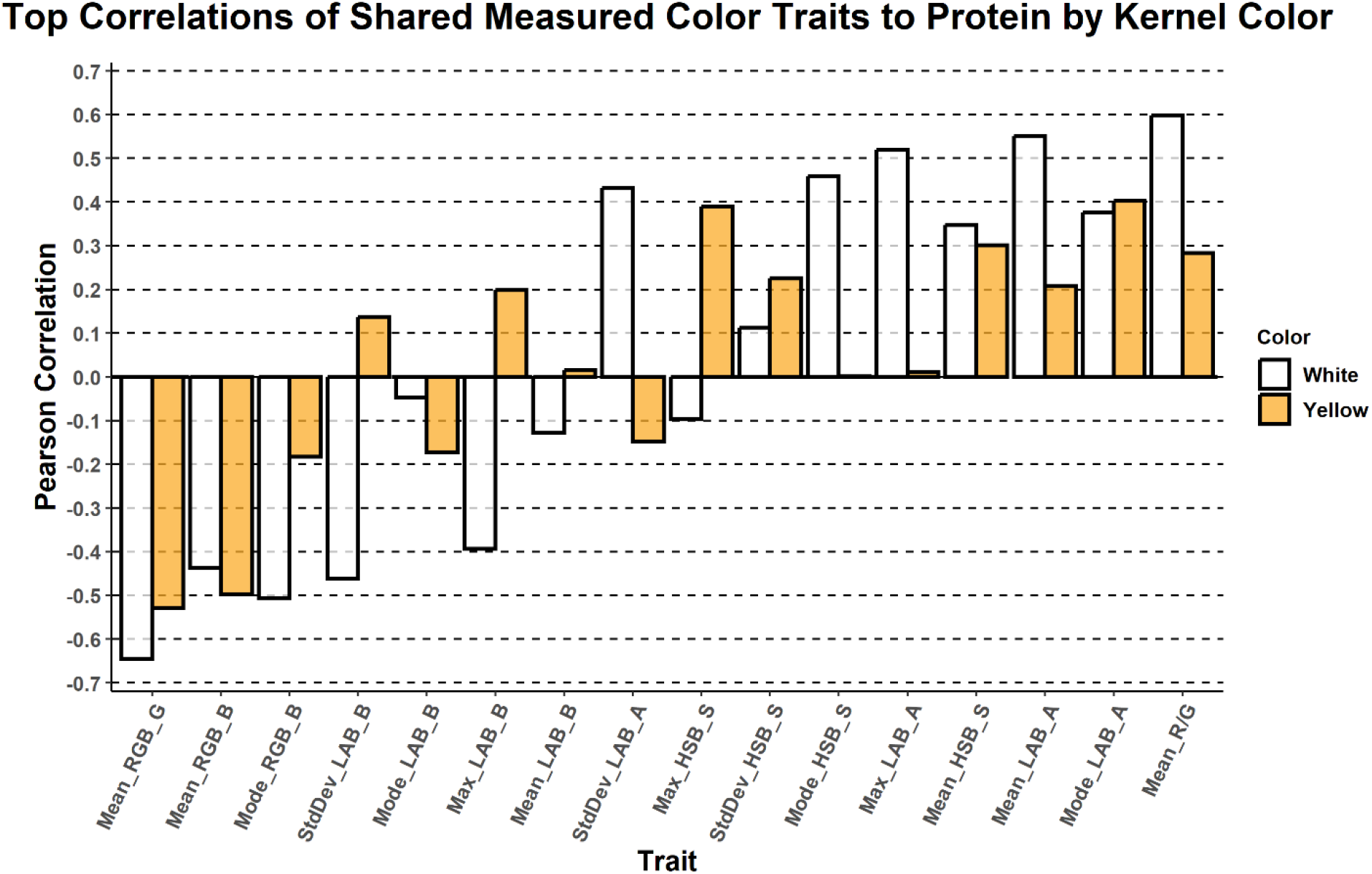
The correlations of the color descriptors that were most robust (r > 0.85) between digital and smartphone cameras with total grain protein concentration measured by NIR, separated by kernel endosperm color.

Moderate correlations between individual color features and grain protein concentration were observed with both camera types, but relationships varied by endosperm color (Supplementary Table 1). No single metric performed consistently across cameras and color classes. As an example, the standard deviation of the RGB Green channel had the strongest correlation of any single feature to NIR grain protein concentration (*r* = -0.74) in the yellow endosperm with smartphone camera class yet showed no meaningful correlation in the white endosperm with smartphone camera (*r* = -0.05). The same trait showed moderate to low correlations in the digital camera data for both yellow (*r* = - 0.59) and (*r* = -0.34) white endosperm. These observations suggest that a subset of color metrics can be reliably transferred between camera types and supports the feasibility of smartphone-based phenotyping of the *Fl2*-RFP reporter. However, it also highlights the importance of stratifying analyses by endosperm color and camera type to reduce confounding factors.

**Table 1.** Color features selected by elastic net regression indicated by (X)

|  | RGB_R | RGB_G | RGB_B | LAB_L | LAB_A | LAB_B | HSB_H | HSB_S | HSB_B |
| --- | --- | --- | --- | --- | --- | --- | --- | --- | --- |
| Mean |  | X |  | X | X |  | X |  |  |
| Max | X |  |  |  | X | X | X |  | X |
| Min |  | X |  | X |  |  |  |  |  |
| Mode | X | X | X | X | X |  | X |  | X |
| StDev |  | X | X |  |  |  |  | X | X |

### 3.4 Prediction of grain protein concentration using *Fl2-RFP*

As it was not immediately clear which features were most informative of the *Fl2*-RFP phenotype, we implemented a modeling approach to select features and generate predictions. Results are shown in Figure 6. Beginning with the set of 52 features extracted from smartphone images, an elastic net regression model selected 27 predictors, reducing the original set by approximately half. It also selected the dummy endosperm color variable, once again underscoring the importance of this effect. The selected predictors included four of five shape metrics; area, aspect ratio, roundness, and solidity, indicating that kernel shape information contributes additional predictive power for grain protein concentration beyond color descriptors. Indeed, when the same modeling approach was employed to predict kernel protein content (concentration multiplied by kernel weight, which is highly proportional to kernel size), the model fit increased even further for both white and yellow kernels. Table 1 displays which color features were selected. Each of the 9 color channels and 5 mathematical summaries are represented at least once, with the RGB Green channel appearing most frequently at 4 times, and mode-based measurements appearing most frequently at 7 times. Although the *Fl2*-RFP reporter is named for its red fluorescence, our analysis suggests the reporter also has an effect on reducing greenness rather than simply increasing redness. In the future, a larger dataset homogenous for endosperm color will help narrow down the feature set that can be used to best describe the phenotype.

**Figure 6.**
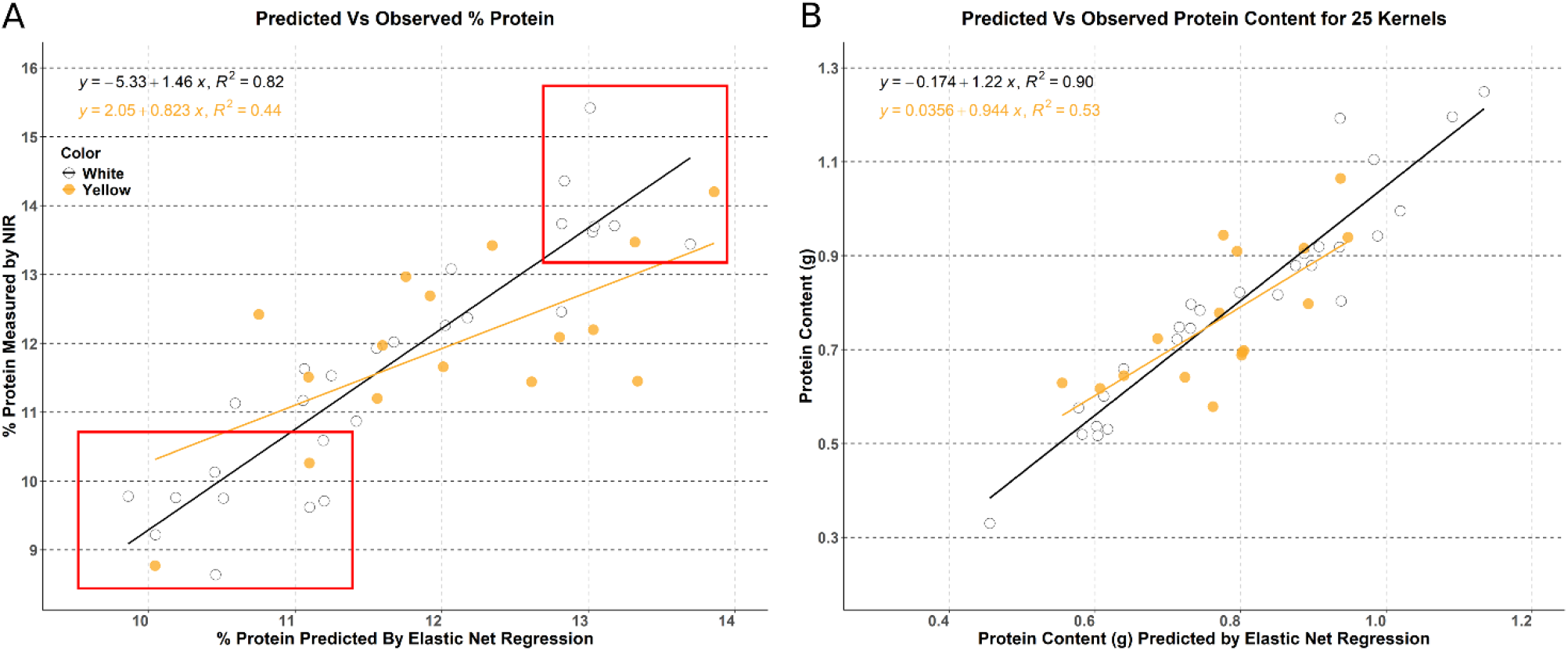
**(A)** Comparison of protein concentration predicted by elastic net regression of imaged *Fl2*-RFP kernels versus that measured by NIR, separated by endosperm color. **(B)** Comparison of protein content for 25 kernels predicted by elastic net regression of imaged *Fl2*-RFP kernels versus calculated by NIR, separated by endosperm color.

The predicted grain concentrations ranged from 9.86% to 13.86%, indicating that the algorithm had difficulty predicting the more extreme values. Overall, the model had an *R*^2^ = 0.68, which can be subdivided into *R*^2^ values of 0.82 for white endosperm and 0.44 for yellow endosperm (Figure 6). However, the RMSE values were similar for all observations (0.93), white endosperm (0.91), or yellow endosperm (0.97). The mean absolute errors (MAE) for the best model (0.76 for all observations) indicates prediction accuracy within one percentage point of protein concentrations measured by NIR. Collectively, these results indicate that maize grain protein concentration can be estimated from smartphone images of kernels expressing the FL2-RFP reporter with sufficient accuracy to effectively select those inbred lines at least 1% lower or 1% higher grain protein concentration than the population average.

## 4 Discussion

Our results demonstrate that the relative intensity of the FL2-RFP reporter can be used to assess α-zein expression, and in some cases overall grain protein concentration in maize kernels. Prior studies in the IPS have shown that selection for total grain protein concentration specifically altered α-zein abundance (Bhattramakki et al., 1996). Accordingly, in both inbreds and hybrids derived from the IPS, there is a high correlation between grain protein concentration, α-zein abundance, and *Fl2*-RFP expression (Lucas et al., 2013). However, this may not be the case for all maize germplasm. Multiple α-zein haplotypes exist among maize inbreds, and these haplotypes differ in both expression level of the α-zeins and in the transcription factors that regulate them (Song & Messing, 2003; Hurst et al., 2021). Given that the α-zein gene family is large, complex, and highly duplicated, the *Fl2*-RFP reporter provides a proxy for rapidly estimating α-zein expression, even when the underlying genomic architecture is difficult to resolve directly.

Kernel protein concentration in maize closely follows that of the maternal genotype (East and Jones, 1920; Letchworth and Lambert 1998). Therefore, the *Fl2*-RFP reporter can be transmitted through a pollen donor and reveal the phenotype of the ear it pollinates (Fig. S1). At least three possible mechanisms have been proposed for the strong maternal effects on grain protein concentration (Moose et al. 2004), including endosperm dosage effects, maternal imprinting of α-zein gene expression, and nutrient supply from vegetative source tissues. Endosperm dosage seems to have minimal impact, because the *Fl2*-RFP phenotype is remarkably consistent among all kernels within hybrid ears (Figure 2A), despite individual kernels varying from 1 to 3 doses of the *Fl2*-RFP transgene in the endosperm. This result agrees with findings from Reggiani et al. (1985) that did not find a significant impact of endosperm dosage on zein concentration in ears from reciprocal crosses between IHP and ILP. Imprinting is an epigenetic mechanism where gene expression is active only when transmitted through one (typically maternal) parent. The expression of some α-zein genes has been observed to show maternal imprinting via DNA demethylation specifically in endosperm tissue (Lund et al., 1995). However, reciprocal crosses involving *Fl2*-RFP illustrate that regardless of genetic background, it is expressed similarly whether transmitted through either the male or female parent. The lack of support for either the endosperm dosage or maternal imprinting hypotheses leaves nutrient supply from vegetative source tissues as most important for maternal control of zein gene expression and seed protein concentration. The demonstration that *Fl2*-RFP accumulation can be modulated by nitrogen supply (Fig. 3) offers direct evidence for this mechanism.

We found that additional pigments in the kernels can alter how *Fl2*-RFP is perceived, such that it is best to restrict measurements to a single endosperm color type when possible. Carotenoids in yellow kernels shift the appearance of kernels containing the *Fl2*-RFP reporter toward orange or red hues, while white kernels display pink tones, and anthocyanin-rich kernels can obscure the signal entirely. To further complicate interpretability, all these pigment classes can occur on the same ear. Even within a color class there is variation in pigment level, as was observed with different shades of yellow endosperm in our inbred panel. This pigment variance likely contributes to unexplained model error in the yellow endosperm class, as it is difficult to disentangle the effects of the *Fl2*-RFP reporter phenotype from the variation in native endosperm color. Maize carotenoids strongly absorb blue light of 460-480nm (Vierstra and Poff, 1981), and thus could impact transmission or reflectance of spectral features captured by cameras. Indeed, Figure 5 shows that the correlation between the Mode RGB Blue feature and NIR protein concentration was reduced in yellow compared to white kernels. Similarly, the color of carotenoids overlaps with green excitation wavelengths for RFP, where abundant RFP may also sufficiently filter light to be detected as a reduction in “greenness”. The need to manually separate kernels for either the *Fl2*-RFP reporter or endosperm color is the most time-consuming portion of our phenotyping, especially when *Fl2*-RFP expression is low. Warman et al. (2021) were able to use computer vision to reliably identify kernels containing fluorescent reporters and anthocyanin pigmentation in images. It may be possible to apply a similar computer vision approach with fluorescence imaging to assist in identifying *Fl2*-RFP kernels.

Several practical considerations emerged from this work. The *Fl2*-RFP reporter is prone to photobleaching, requiring careful handling of seed material during development and storage to preserve the phenotype until it is imaged. Furthermore, our analysis revealed that not all image-derived features are equally stable across imaging conditions, even while taking precautions to minimize differences. We captured images in a completely dark room with a consistent artificial light source, and matched device settings as closely as possible. The smartphone offered fewer manual controls than the digital camera, which reduced cross-device reproducibility. Despite these discrepancies, many features derived from smartphone images yielded equivalent information quality to enable robust trait predictions. Shape descriptors were highly reproducible across both digital and smartphone cameras and provided additional valuable information about kernel growth that enhanced prediction of grain protein concentration. Only a subset of color features showed strong cross-device concordance, indicating that imaging hardware can influence how specific aspects of the phenotype are captured. These results highlight the need for standardized acquisition protocols and, when possible, the use of a single camera type or preset settings for comparative analyses. Nonetheless, the strong agreement observed for several color traits supports the feasibility of smartphone-based phenotyping for *Fl2*-RFP expression.

The *Fl2*-RFP reporter offers a new approach to easily monitor α-zein accumulation in maize genetics experiments and breeding improvement. Since the initial discovery that the *fl2* mutation enhances amino acid balance of maize grain by reducing α-zein (Nelson et al, 1965; Coleman et al., 1997), the “zein reduction” phenotype has been targeted by a variety of molecular breeding strategies (e.g. Huang et al., 2006) to improve nutritional quality for food and feed, such as the development of Quality Protein Maize (QPM, reviewed by Prasanna et al., 2001). Like many such efforts, direct use of *fl2* was discontinued for QPM due to undesirable effects of low α-zein on kernel texture. The *Fl2*-RFP reporter may enable more effective selection for reducing α-zein while maintaining kernel density and hardness, a phenotype evident in the B73 x IRHP hybrid (Figure 2A).

Although only one gene among the many contributing to grain protein concentration in maize, easy measurement of FL2-RFP and its functional restriction to nitrogen storage makes it a useful biosensor for tracking the interactive effects of variation in both hybrid genotype and nitrogen supply in programming crop yield, nutritional quality and sustainability. We have already demonstrated the capacity to separate N-responsiveness of α-zeins from total protein at a research scale (Figure 2B), such experiments could be readily scaled to larger populations through crossing to common FL2-RFP testers, or inclusion of only a small proportion of FL2-RFP hybrid plants as pollen donors for broader field-level reporting.

Higher nitrogen fertilizer applications are known to increase grain protein concentration and some studies have also documented higher α-zein accumulation. However, historical corn breeding for improved yields, yield N response, and N use efficiency has also reduced grain protein concentration (Haegele et al. 2013; DeBruin et al., 2017; King et al; 2024), despite these hybrids typically being grown with at least sufficient and often excessive nitrogen needed to maximize grain yield (Baum et al., 2025). The observation that N fertilization increases yield but not protein concentration in recent hybrids suggests that breeding may have reduced the N-responsiveness of α-zeins. Dual phenotyping with NIR and *Fl2*-RFP reveals genetic variation exists for this trait (Figure 2B) and suggests strategies for increased selection accuracy and accelerated genetic gain to reduce α-zeins and overall seed nitrogen accumulation. Such phenotypes are desirable for efforts to redesign maize for more sustainable cropping systems (Odilón Ojeda-Rivera et al., 2025). Importantly, the *Fl2*-RFP reporter can be included during the breeding program and then removed before commercialization to avoid potential regulatory issues with transgenic maize.

## 5 Conclusion

We validated that the *Fl2*-RFP reporter can be visualized at many different scales, with its most practical advantage being clearly detectable under white light, eliminating the need for specialized imaging equipment. A standard smartphone camera can effectively document the *Fl2*-RFP reporter phenotype, which can then be quantified through an automated imaging pipeline, providing a rapid, repeatable, and scalable method for investigating α-zein expression in maize kernels. With further standardization, this reporter system and workflow can be broadly applied to monitor genotype-by-nitrogen interactions within maize seeds, which will enhance genetic dissection and selection for optimizing this phenotype in breeding programs.

## 6 Conflict of Interest

The authors declare that the research was conducted in the absence of any commercial or financial relationships that could be construed as a potential conflict of interest.

## 7 Author Contributions

CL: formal analysis, investigation, methodology, software, visualization and writing; NJH: Conceptualization, Investigation, Visualization; Writing – original draft; CT: Conceptualization, Investigation, Visualization; Writing – review and editing; SPM: Conceptualization, Investigation, Visualization, Funding acquisition, Supervision, Writing – review and editing.

## 8 Funding

Funding for this project was provided by the Illinois Maize Breeding and Genetics Laboratory, the Denton Eugene and Elizabeth Alexander Professorship in Maize Breeding and Genetics (to SPM), and graduate fellowships from Illinois Corn to CL, NJH, CT. In addition, early stages of the research were supported by USDA-NIFA award number 2011-67013-30041, NSF Grant No. DBI-2019674, and sponsorship by the National Science Foundation (award IOS-102594) of NH participation at the NUPRIME workshop for Chromatin Structure and Genome Response in Maize.

## Acknowledgments

We are grateful to David Jackson and Anne Sylvester for providing original seeds of Fl2-RFP transgenic lines. We acknowledge Jessica Bubert for her help in producing Fl2-RFP hybrid ears from nitrogen-responsive field trials. We also acknowledge the special contributions of Jarai Carter and Samantha Shaffer for assistance in capturing and data processing of camera images.

## Notes

### Competing Interest Statement

The authors have declared no competing interest.

## References

Bass, H.W., Wear, E.E., Lee, T., Hoffman, G.G., Gumber, H.K., Allen, G.C., et al. (2014). A maize root tip system to study DNA replication programmes in somatic and endocycling nuclei during plant development. Journal of experimental botany, 65(10), 2747–2756. doi: 10.1093/jxb/ert470

Baum, M.E., Sawyer, J.E., Nafziger, E.D., Castellano, M.J., McDaniel, M.D., Licht, M.A., et al. (2025). The optimum nitrogen fertilizer rate for maize in the US Midwest is increasing. Nature Communications, 16(1), p.404. doi: 10.1038/s41467-024-55314-7

Bhattramakki, D., Sachs, M.M., and Kriz, A.L. (1996). Expression of genes encoding globulin and prolamin storage proteins in kernels of Illinois long term chemical selection strains. Crop Sci 36, 1029–1036. doi: 10.2135/cropsci1996.0011183X003600040036x

Campbell, R.E., Tour, O., Palmer, A.E., Steinbach, P.A., Baird, G.S., Zacharias, D.A., et al. (2002). A monomeric red fluorescent protein. PNAS. doi: 10.1073/pnas.082243699

Coleman, C.E., Clore, A.M., Ranch, J.P., Higgins, R., Lopes, M.A., & Larkins, B.A. (1997). Expression of a mutant α-zein creates the floury 2 phenotype in transgenic maize. PNAS, 94(13), 7094–7097.

Coleman, C.E. and Larkins, B.A. (1999). “The prolamins of maize.” In Seed proteins (pp. 109–139). Dordrecht: Springer Netherlands.

DeBruin, J.L., Schussler, J.R., Mo, H. and Cooper, M. (2017). Grain yield and nitrogen accumulation in maize hybrids released during 1934 to 2013 in the US Midwest. Crop Sci. 57, 1431–1446.

Dudley, J.W., and Lambert, R.J. (2004). “100 Generations of Selection for Oil and Protein in Corn.” In Plant Breeding Reviews, (Wiley), 79–110. doi: 10.1002/9780470650240.ch5

East, E.M. and Jones, D.F. (1920). Genetic studies on the protein content of maize. Genetics, 5(6), p.543.

Feng, L., Zhu, J., Wang, G., Tang, Y., Chen, H., Jin, W., et al. (2009). Expressional profiling study revealed unique expressional patterns and dramatic expressional divergence of maize α-zein super gene family. Plant Mol Biol 69, 649–659. doi: 10.1007/s11103-008-9444-z

Flint-Garcia, S.A., Bodnar, A.L., and Scott, M.P. (2009). Wide variability in kernel composition, seed characteristics, and zein profiles among diverse maize inbreds, landraces, and teosinte. Theoretical and Applied Genetics 119, 1129–1142. doi: 10.1007/s00122-009-1115-1

Haegele, J.W., Cook, K.A., Nichols, D.M., & Below, F.E. (2013). Changes in nitrogen use traits associated with genetic improvement for grain yield of maize hybrids released in different decades. Crop Science, 53(4), 1256–1268.

Hopkins, C.G. (1899). Improvement in the chemical composition of the corn kernel. Journal of the American Chemical Society, 21(11), 1039–1057.

Hopkins, C.G., Smith L.H., and East E.M. (1903). The structure of the corn kernel and the composition of its different parts. Ill. Agr. Exp. Sta. Bul, 87, p.I903.

Huang, S., Frizzi, A., Florida, C.A., Kruger, D.E. and Luethy, M.H. (2006). High lysine and high tryptophan transgenic maize resulting from the reduction of both 19-and 22-kD α-zeins. Plant molecular biology, 61(3), pp.525–535.

Hunter, B.G., Beatty, M.K., Singletary, G.W., Hamaker, B.R., Dilkes, B.P., Larkins, B.A., et al. (2002). Maize Opaque Endosperm Mutations Create Extensive Changes in Patterns of Gene Expression. Plant Cell 14, 2591–2612. doi: 10.1105/tpc.003905

Hurst, P., Schnable, J.C., and Holding, D.R. (2021). Tandem duplicate expression patterns are conserved between maize haplotypes of the α-zein gene family. Plant Direct 5. doi: 10.1002/pld3.346

Kim, C.S., Woo, Y.M., Clore, A.M., Burnett, R.J., Carneiro, N.P. and Larkins, B.A., 2002. Zein protein interactions, rather than the asymmetric distribution of zein mRNAs on endoplasmic reticulum membranes, influence protein body formation in maize endosperm. The Plant Cell, 14(3), pp.655–672. doi: 10.1105/tpc.010431

King, K., Ferela, A., Vyn, T.J., Trifunovic, S., Eudy, D., Hurburgh, C., Lamkey, K.R., & Archontoulis, S.V. (2024). Genetic gains in short-season corn hybrids: Grain yield, yield components, and grain quality traits. Crop Science, 64, 710–725.

Krishnakumar, V., Choi, Y., Beck, E., Wu, Q., Luo, A., Sylvester, A., et al. (2015). A maize database resource that captures tissue-specific and subcellular-localized gene expression, via fluorescent tags and confocal imaging (maize cell genomics database). Plant Cell Physiol 56, e12. doi: 10.1093/pcp/pcu178

Lending, C.R., and Larkins, B.A. (1989). Changes in the Zein Composition of Protein Bodies during Maize Endosperm Development. Plant Cell 1, 1011–1023.

Letchworth, M.B., and Lambert, R.J. (1998). Pollen parent effects on oil, protein, and starch concentration in maize kernels. Crop Sci 38, 363–367. doi: 10.2135/cropsci1998.0011183X003800020015x

Lopes, M.A., and Larkins, B.A. (1993). Endosperm Origin, Development, and Function. Plant Cell 5, 1383–1399.

Lucas, C.J., Zhao, H., Schneerman, M., and Moose, S.P. (2013). “Genomic Changes in Response to 110 Cycles of Selection for Seed Protein and Oil Concentration in Maize.” In Seed Genomics, (Wiley-Blackwell), 217–236. doi: 10.1002/9781118525524.ch12

Lund, G., Ciceri, P. and Viotti, A. (1995), Maternal-specific demethylation and expression of specific alleles of zein genes in the endosperm of Zea mays L.. The Plant Journal, 8: 571–581.

Nelson, O.E., Mertz, E.T., & Bates, L.S. (1965). Second mutant gene affecting the amino acid pattern of maize endosperm proteins. Science, 150(3702), 1469–1470.

Makanza, R., Zaman-Allah, M., Cairns, J.E., Eyre, J., Burgueño, J., Pacheco, Á., et al. (2018). High-throughput method for ear phenotyping and kernel weight estimation in maize using ear digital imaging. Plant Methods 14. doi: 10.1186/s13007-018-0317-4

Miller, N.D., Haase, N.J., Lee, J., Kaeppler, S.M., de Leon, N., and Spalding, E.P. (2017). A robust, high-throughput method for computing maize ear, cob, and kernel attributes automatically from images. Plant Journal 89, 169–178. doi: 10.1111/tpj.13320

Mohanty, A., Luo, A., DeBlasio, S., Ling, X., Yang, Y., Tuthill, D.E., et al. (2009). Advancing cell biology and functional genomics in maize using fluorescent protein-tagged lines. Plant Physiology 149, 601–605. doi: 10.1104/pp.108.130146

Moose, S.P., Dudley, J.W., and Rocheford, T.R. (2004). Maize selection passes the century mark: A unique resource for 21st century genomics. Trends Plant Sci 9, 358–364. doi: 10.1016/j.tplants.2004.05.005

Ojeda-Rivera, J.O., Barnes, A.C., Ainsworth, E.A., Angelovici, R., Basso, B., Brindisi, L.J., et al. (2025). Designing a nitrogen-efficient cold-tolerant maize for modern agricultural systems. The Plant Cell, 37(7), p.koaf139.

Paulis, J.W. and Wall, J.S., (1977). Comparison of the protein compositions of selected corns and their wild relatives, teosinte and Tripsacum. Journal of Agricultural and Food Chemistry, 25(2), pp.265–270.

Prasanna, B.M., Vasal, S.K., Kassahun, B. and Singh, N.N., 2001. Quality protein maize. Current science, pp. 1308–1319.

R Core Team (2026). R: A Language and Environment for Statistical Computing. R Foundation for Statistical Computing, Vienna, Austria. https://www.R-project.org/.

Reggiani, R., Soave, C., Di Fonzo, N., Gentinetta, E. and Salamini, F. (1985). Factors affecting starch and protein content in developing endosperms of high and low protein strains of maize (Zea mays L.). Genetica Agraria 39:221–232.

Schindelin, J., Arganda-Carreras, I., Frise, E., Kaynig, V., Longair, M., Pietzsch, T., et al. (2012). Fiji: An open-source platform for biological-image analysis. Nat Methods 9, 676–682. doi: 10.1038/nmeth.2019

Song, R. and Messing, J. (2003). Gene expression of a gene family in maize based on noncollinear haplotypes. PNAS, 100(15), pp.9055-9060. doi: 10.1073/pnas.1032999100

Strock, C. (2021). Protocol for extracting basic color metrics from Images in ImageJ/Fiji. Zenodo. doi:10.5281/zenodo.5595203

Tay J.K., Narasimhan B., Hastie T. (2023). “Elastic Net Regularization Paths for All Generalized Linear Models.” Journal of Statistical Software, 106(1), 1–31. doi:10.18637/jss.v106.i01.

Tollenaar, M., and Dwyer, M. (1999). “Physiology of Maize,” in Crop Yield, Physiology and Processes, eds. D. L. Smith and C. Hamel.

Tsai, C.L., I. Dweikat and C.Y. Tsai. (1990). Effects of source supply and sink demand on the carbon and nitrogen ratio in maize kernels. Maydica. 35: 391–397.

Uribelarrea, M., Below, F.E., and Moose, S.P. (2004). Grain Composition and Productivity of Maize Hybrids Derived from the Illinois Protein Strains in Response to Variable Nitrogen Supply. Crop Sci 44, 1593–1600. doi: 10.2135/cropsci2004.1593

Vierstra, R.D. and Poff, K.L. (1981). Role of carotenoids in the phototropic response of corn seedlings. Plant Physiology, 68(4), pp.798–801.

Warman, C., Sullivan, C.M., Preece, J., Buchanan, M.E., Vejlupkova, Z., Jaiswal, P., et al. (2021). A cost-effective maize ear phenotyping platform enables rapid categorization and quantification of kernels. Plant Journal 106, 566–579. doi: 10.1111/tpj.15166

Wenck, A., Pugieux, C., Turner, M., Dunn, M., Stacy, C., Tiozzo, A., et al. (2003). Reef-coral proteins as visual, non-destructive reporters for plant transformation. Plant Cell Rep 22, 244–251. doi: 10.1007/s00299-003-0690-x

Woo, Y.-M., Wang, D., Hu, N., Larkins, B.A., and Jung, R. (2001). Genomics Analysis of Genes Expressed in Maize Endosperm Identifies Novel Seed Proteins and Clarifies Patterns of Zein Gene Expression. Plant Cell 13, 2297–2317. doi: 10.1105/tpc.010240

Wu, Q., Luo, A., Zadrozny, T., Sylvester, A. and Jackson, D. (2013). Fluorescent protein marker lines in maize: generation and applications. International Journal of Developmental Biology, 57(6-7-8), pp. 535–543.

Zou, H. and Hastie, T. (2005). Regularization and variable selection via the elastic net. Journal of the Royal Statistical Society Series B: Statistical Methodology, 67(2), pp.301–320.

