## Supplementary material for "Rapid, Non-Destructive Visualization of α-Zein Expression and Grain Protein Concentration in Maize Using the *Floury2*-RFP Reporter Transgene": Figure S1

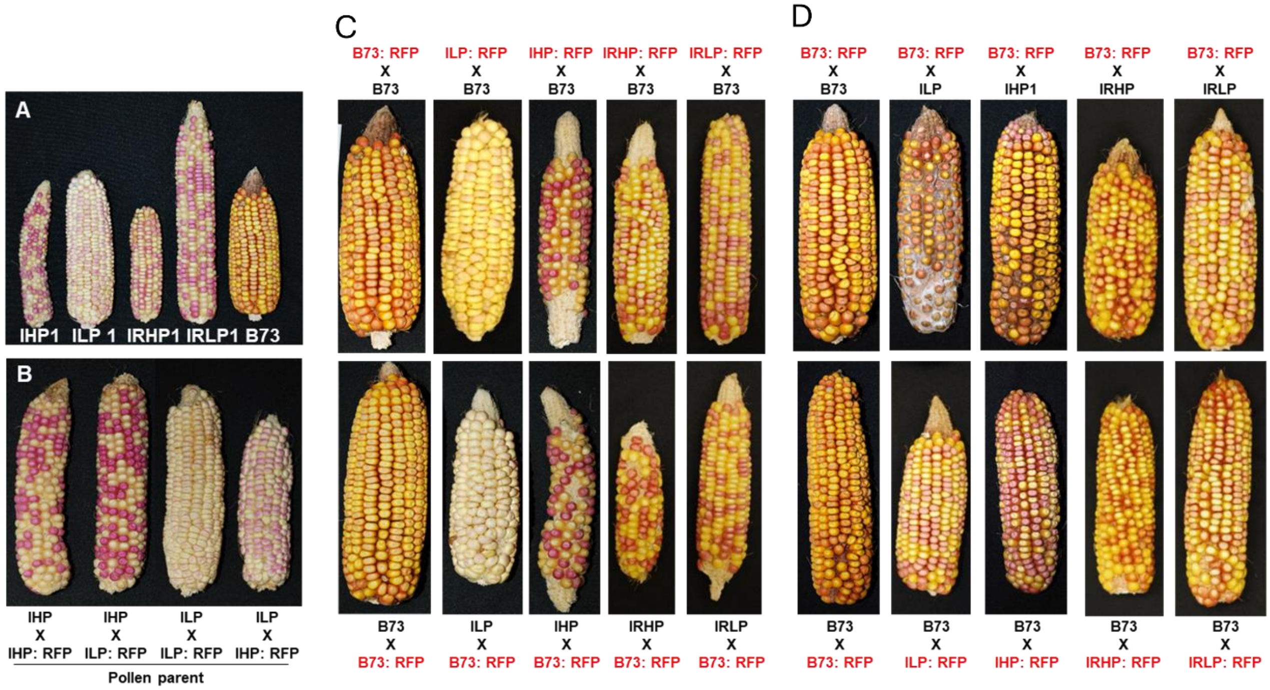


**Supplementary Figure 1. Images of ears expressing Fl2-RFP in the IPS and B73 inbred backgrounds.** **(A)** Ears from the introgression lines heterozygous for Fl2-RFP, where the maternal parent carries the Fl2-RFP transgene. IHP1 = Illinois High Protein1, ILP1 = Illinois Low Protein1, IRHP1 = Illinois Reverse High Protein1, IRLP1 = Illinois Reverse Low Protein1. **(B)** Ears from reciprocal crosses of the same IHP1 and ILP1 parents shown in (A), but where the paternal parent transmits Fl2-RFP. **(C)** Ears from reciprocal crosses of the inbreds shown in (A), where the maternal genetic background was varied in combination with a B73 male. **(D)** Ears from reciprocal crosses of the inbreds shown in panel (A), where different male parents transmitted the Fl2-RFP transgene to B73 ears.
